# OpenEHR modeling for genomics in clinical practice

**DOI:** 10.1101/194720

**Authors:** Cecilia Mascia, Paolo Uva, Simone Leo, Gianluigi Zanetti

## Abstract

The increasing usage of high throughput sequencing in personalized medicine brings new challenges to the realm of healthcare informatics. Patient records need to accommodate data of unprecedented size and complexity as well as keep track of their production process. In this work we present a solution for integrating genomic data into electronic health records via openEHR archetypes. We introduce new genomics-specific archetypes based on the popular variant call format and show their applicability to a practical use case. Finally, we discuss their structure in comparison with the HL7^®^ FHIR^®^ standard.

## 1 Introduction

Next generation sequencing (NGS) techniques have recently enabled a substantial advancement in the treatment of genetic disorders, paving the way for personalized therapies based on each patient’s specific set of genomic variations. Whole-exome (WES) and whole-genome sequencing (WGS) have become increasingly common in the past years, thanks mainly to the progress and cost reduction in high-throughput sequencing technologies^1,2^. This trend suggests that genomic data are going to play an increasingly important role within medical practice in the near future.

Entities related to a patient’s clinical history can be modeled according to a spoke-hub paradigm, where the different actors (such as clinicians, specialists, hospitals, labs and the patient itself) all revolve around the Electronic Health Record (EHR), the central collection point of all clinical data provided by each actor. Due to its high relevance in modern healthcare, genomic information should be a fundamental part of EHRs. In practice, however, most current EHR systems are not capable of handling such data ^1,2,3,4^, since their sheer size and complexity entail new technological challenges in terms of storage, integration, visualization, computational analysis, querying and presentation. Moreover, where supported, genomic information is often made available as unformatted plain text, whereas a structured, machine-readable format would greatly enhance its shareability and reusability. Finally, particular care must be taken to ensure interoperability with other medical informatics standards. These practical difficulties severely hinder the translation of genetic information into concrete clinical actions.

In this work we present a new openEHR^5^ model for genomic data produced in the context of sequence variation analysis, specifically focused on machine readability and computability. We show its applicability to a concrete use case and its interoperability with existing clinical standards. Finally, we discuss our approach in comparison with related work.

### 1.1 OpenEHR specifications

OpenEHR is an open standard for health data that focuses on the semantic interoperability between EHR systems. It adopts a multi-level modeling approach based upon the Reference Model (RM), a set of classes that represent logical EHR structures and demographic data. The next level consists of a library of *archetypes*, reusable models that define a maximal set of attributes related to a particular subject. Archetypes, in turn, can be combined into *templates*, hierarchical context-specific data sets. Archetypes and templates undergo an iterative web-based review process, at the end of which they are published in the official openEHR repository, the Clinical Knowledge Manager (CKM)^6^, and made available for clinical use.

Archetypes belong to four main groups: *compositions*, which represent commonly used clinical documents; *sections*, corresponding to document headings; *entries*, the most common and fundamental building blocks of an EHR (further divided into *observations*, *evaluations*, *instructions* and *actions*); *clusters*, reusable sub-structures that allow elements to be grouped and repeated. The model presented here makes use of the observation (data obtained by a direct observation or measurement) and cluster archetype classes.

Due to the amount of domain-specific expertise required, EHR systems development needs to be carried out in close cooperation with clinical stakeholders. The main challenges in this process are establishing a common ground for communication between technicians and clinicians and addressing the frequent advancement of clinical knowledge. In the case of genomic content, this is further exacerbated by the size and structure issues discussed above. The multi-level openEHR approach helps mitigate these problems by separating structure and content: only the first modeling level (RM classes) is implemented in software, while formal definitions of clinical content (archetypes and templates) are external. This means that EHR repositories can be developed independently from the content they will store. Moreover, EHR systems can be kept small, maintainable and self-adaptable to archetypes and templates that may be developed in the future^7^. Finally, this decoupling facilitates contributions by non-technical professionals, who can formalize their clinical knowledge via user-friendly modeling tools.

### 1.2 Structured approach

Integrating NGS data into EHRs is hard due to two main reasons^3^:

- their size (up to hundreds of gigabytes) and complexity (e.g., irregular degree of nesting in laboratory test outputs);
- the way they are generated and manipulated, with processing pipelines where each step depends on a multitude of software parameters and resource databases.

Currently available EHR systems typically collect three types of data: granular (e.g., laboratory test results), text (e.g., pathology reports) and media (e.g., medical imaging). As mentioned earlier, due to the lack of specific data structures genomic data is often included in text form, which makes their analysis and reuse particularly cumbersome. The adoption of a structured format would greatly simplify the processing and transfer of clinical genomic data, effectively enhancing its medical value. The chosen format should enable the efficient management of complex clinical content by organizing it into standard reusable entities. At the same time, it should allow the specification of data semantics, while ensuring that the original meaning is preserved in case of sharing. Finally, it should preserve the data history, with regard to both the operations performed and the auxiliary resources involved in the transformation process (such as external databases, genomic references, etc.). In the remainder of this paper we show how to use the openEHR approach to address these issues.

## 2 Materials and methods

### 2.1 Genomic data

WGS and WES have rapidly changed research on the genetic causes of rare and complex diseases. Independently from the technology used, NGS data analysis can be broken down into two main steps^8^: the *analytic wet bench process* and the *bioinformatic analysis of sequence data* (Fig. 1).

**Figure 1:**
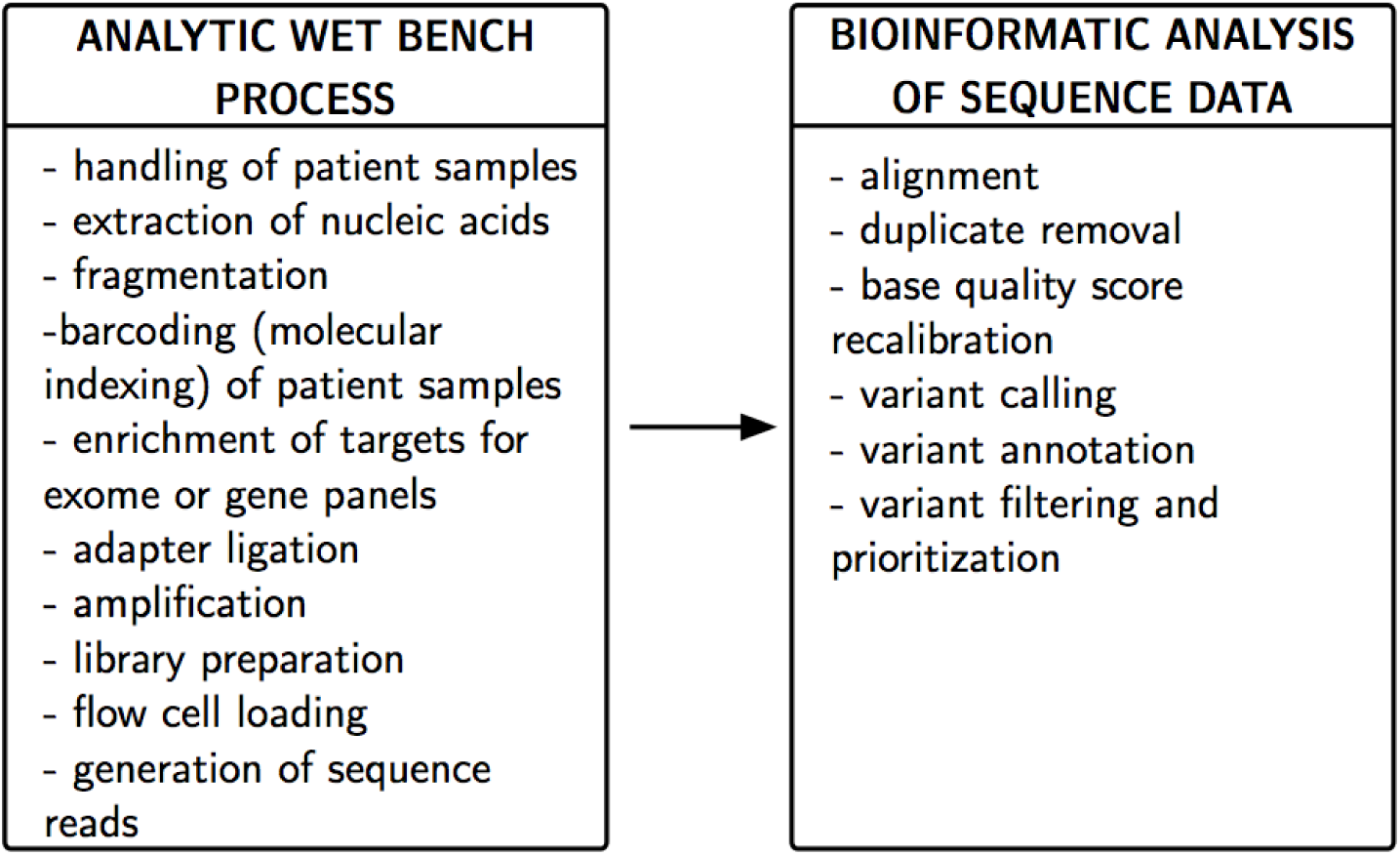
Main NGS steps, grouped accordingly to the College of American Pathologists^8^.

The process starts with DNA extraction and its fragmentation into small pieces. NGS instruments produce billions of shorts sequence reads (strings representing ordered DNA nucleotides in a fragment) simultaneously in a single machine run. Reads are subsequently aligned (mapped) to a reference sequence through dedicated software, a computationally intensive operation due to the large size of the reference. Aligned reads are then processed to correct for technical biases and, finally, genomic variants (the differences between the sample and the reference) are identified and reported, along with additional information such as the coverage depth (average number of reads that align to known reference bases) and accuracy measures. Due to the large amount of variants that can be detected, generating useful results requires one or more filtering steps to prioritize potential disease-causing mutations^9,10,11,12^. This is commonly done by annotating variants with references to public databases such as dbSNP^13^. Common scenarios in human NGS data analysis include the discovery of disease-associated variant(s) for mendelian disorders, the identification of candidate genes responsible for complex diseases and the detection of constitutional and somatic mutations in cancer studies^9^.

Genomic data can be seen as a multilayer structure composed of three levels:

- **Raw data**, the direct output of the sequencing process, consist of millions of short sequence reads, text strings containing the detected DNA nucleotides together with their associated quality scores (statistical predictions of accuracy).
- **Derived data** represents the observed variants, along with additional measurements (e.g., coverage);
- **Annotated data** corresponds to variants filtered according to additional a-priori information, as discussed above.

With respect to Fig. 1, the first level falls within the wet bench block, while the other two are part of the bioinformatic pipeline.

A common bioinformatic workflow for variant identification from NGS data includes the following steps^14^ (Fig. 2):

- **Alignment**: short reads are aligned to the reference to produce a file in SAM/BAM format^15^, sorted by genomic coordinate. Initial mappings are further processed to address technical biases (removal of duplicate sequences, recalibration of base quality scores).
- **Variant calling**: alignment files output by the previous step are processed to detect variations between the sample and the reference. Since variations may also arise from mapping and sequencing artifacts, variant calling software should carefully balance sensitivity to minimize false positives. The output, typically in tabular format, contains the list of variant sites, the individual genotypes and additional information such as coverage and genotyping accuracy.
- **Variant annotation**: assigning functional information to genomic variants is a crucial step in the analysis of sequencing data, enabling researchers to focus on the potential disease-causing variants. Many types of biological information can be associated to variants: the position in transcript sets (e.g., UCSC, RefGene, GENCODE, ENSEMBL), whether the variant is known or novel based on dbSNP, the prediction of their impact on the protein structure and function according to different models (e.g., SIFT, PolyPhen2, LRT, MutationTaster, MutationAssessor, FATHMM), sequence conservation (e.g., PhyloP, PhastCons), known associations of the variant with diseases (e.g., OMIM, ClinVar) and allele frequency in reference populations (e.g., ESP6500, 1000 Genomes Project, gnomAD).

**Figure 2:**
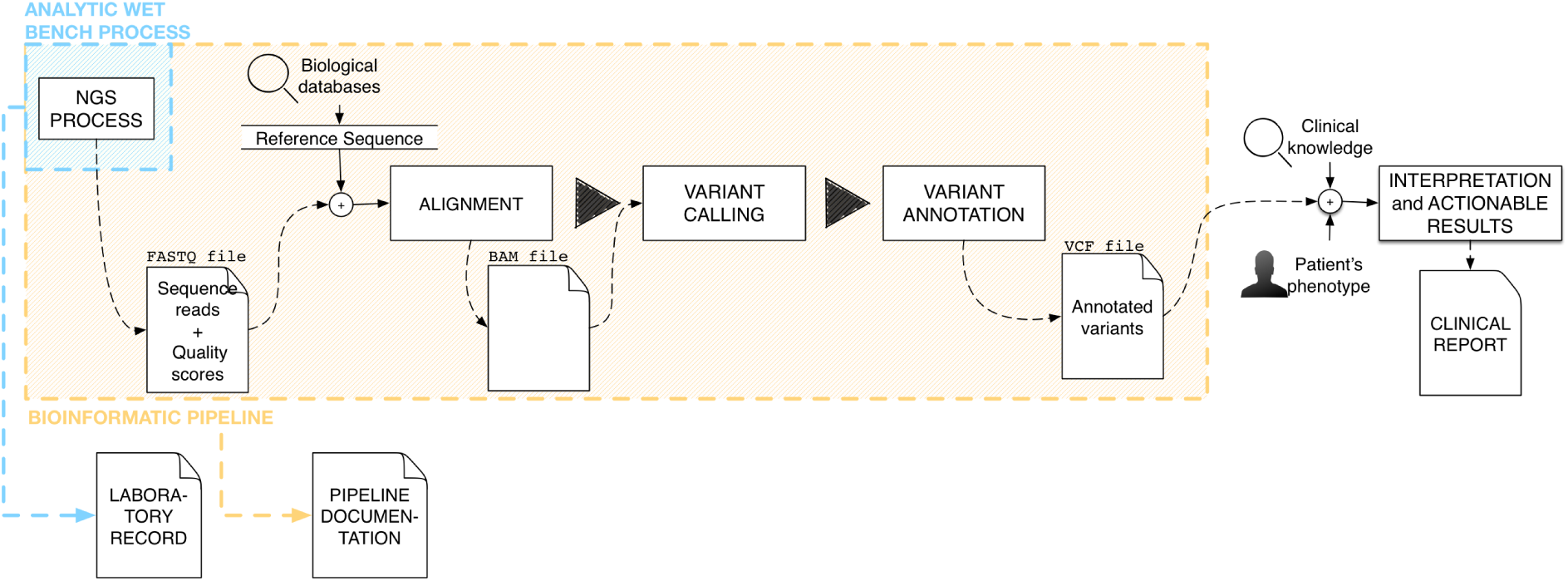
Schematic representation of a bioinformatic pipeline for NGS variant detection.

### 2.2 OpenEHR modeling approach

Figure 3 summarizes the context from a high level perspective: the clinician receives phenotypic and genetic (laboratory block) patient data, and interprets them in combination with prior clinical knowledge and experience to produce actionable results. It should be noted that, before reaching the clinical side, laboratory output should be reorganized to assume a structured form. The openEHR model allows to achieve this through the archetypes formalism.

**Figure 3:**
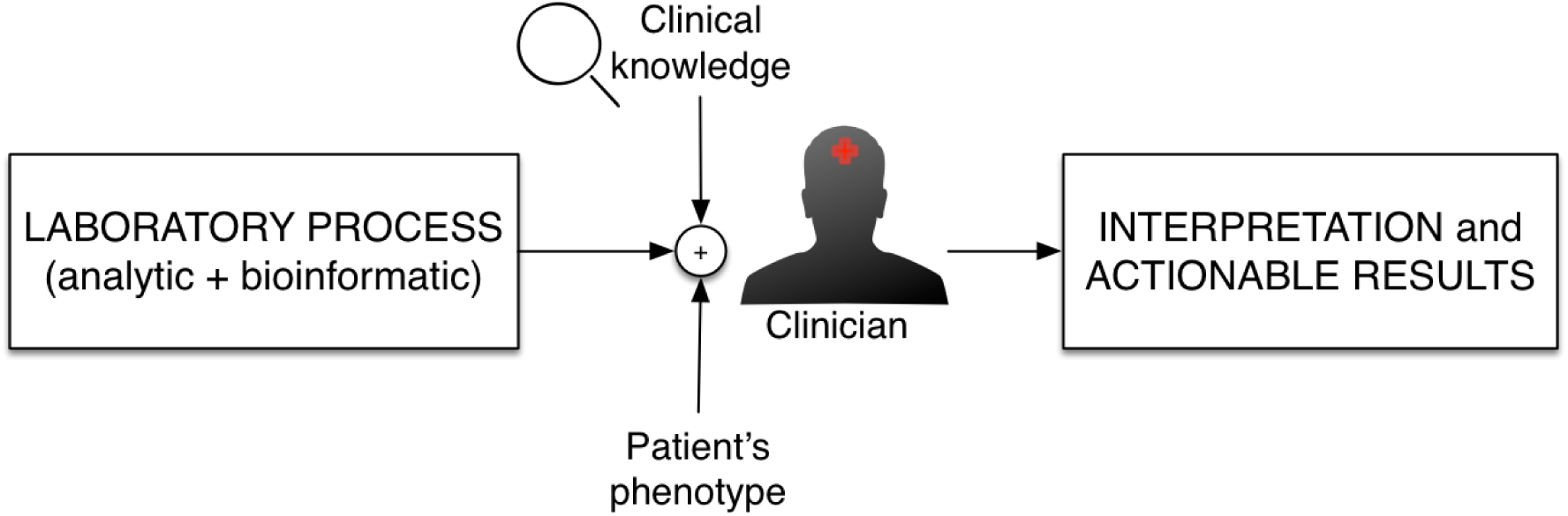
Block diagram of the process leading from data to clinical actions.

As discussed above, genetic data sets are produced via complex pipelines that involve the use of different algorithms, parameters and reference entities. To provide a comprehensive and reusable model for genomic data, archetypes should be able to keep track of all this information in a structured way. Since all of these elements can change over time, to ensure reproducibility we include this information as external references in our model.

Common openEHR modeling practice follows (possibly with recursion) these steps:

- define the data to be represented;
- search model repositories for reusable archetypes and, possibly, map existing archetype nodes with clinical attributes;
- create new archetypes or specialize existing ones for the specific domain being modeled.

In our case, the model has to represent the output of the bioinformatic pipeline, i.e., information about the variants detected at specific positions in the genome. Due to its wide adoption for the representation of genomic data, we have designed our archetypes to mirror the Variant Call Format (VCF)^16^. As shown in Fig. 4, a VCF file consists of meta-information lines, a header line and data lines (body).

**Figure 4:**
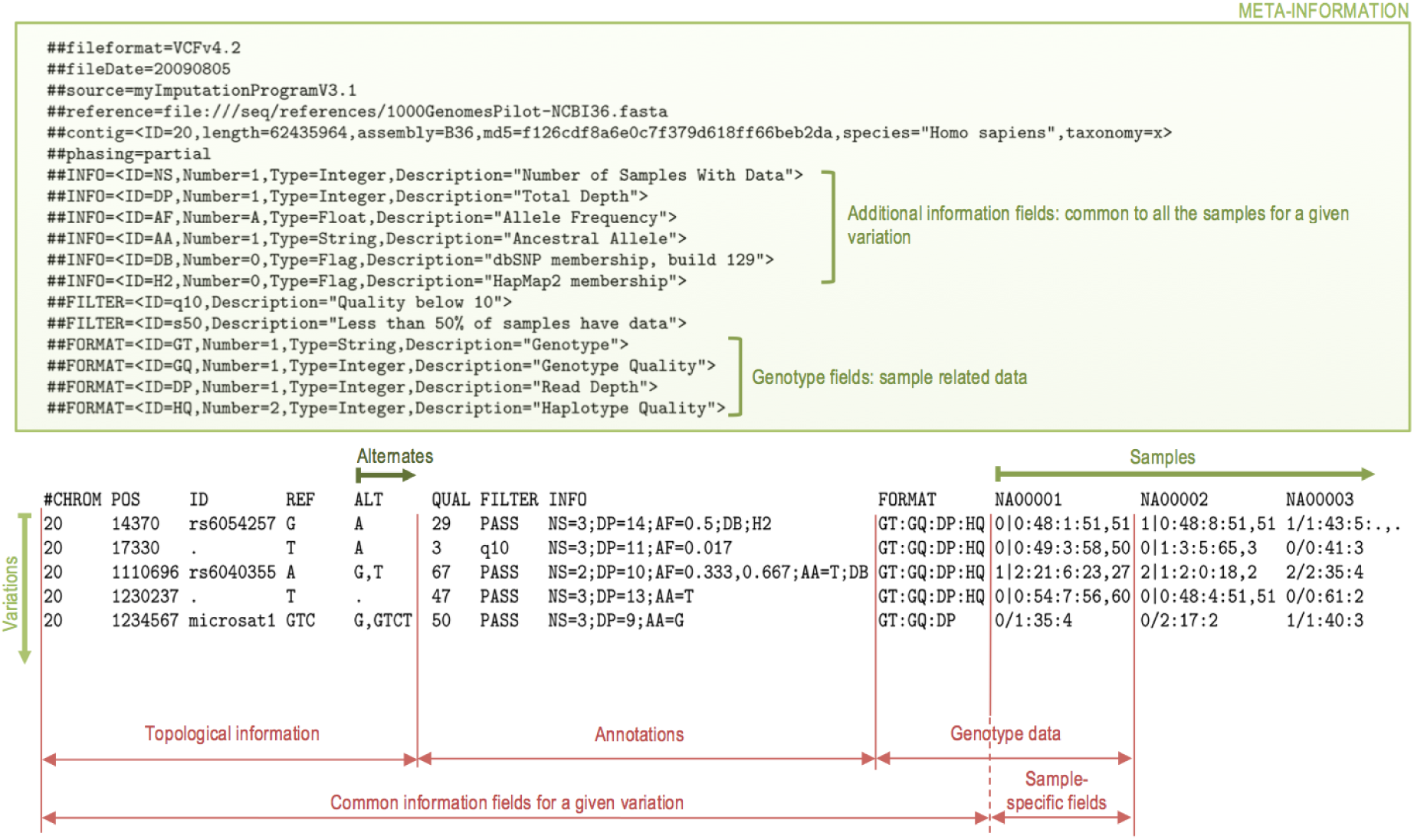
The main section of a typical VCF file. Modified figure from the VCF specifications^16^.

Each row in the VCF body corresponds to a specific location in the genome where a single variation occurs, possibly with more than one alternate allele. All rows start with eight mandatory fields (although one or more could be empty): `#CHROM`, `POS`, `ID`, `REF`, `ALT`, `QUAL`, `FILTER` and `INFO`. If there are multiple alternate alleles called on at least one sample for a given position, these are reported as a comma separated list under the ALT field. The INFO field (see Fig. 5) gives additional information on the observed variant, in a format specified in the meta-information.

**Figure 5:**
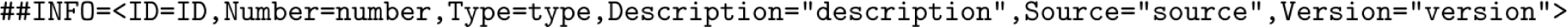
The VCF INFO field^16^.

Sample-specific information (such as coverage or genotype accuracy), is encoded in the FORMAT column, whose structure is also specified in the meta-information block. In addition to the mandatory ones, VCF allows to specify additional fields to accommodate specific domain annotations. The most common ones have already been included in the model, while additional ones can be specified via custom clusters.

As mentioned earlier, before embarking on the creation of new archetypes, a thorough search of existing public domain ones must be performed. At the time of writing, no archetype in the international instance of the CKM^6^ was suitable for genomic data. To fill this gap, we developed a new set of archetypes that are described in Sec. 3. This work has been carried out with the LinkEHR Studio software developed by the Universitat Politècnica de València and VeraTech for Health^17^.

## 3 Results

A few entities involved in the sequencing workflow — such as specimen and device — were already supported in the CKM, while others required the development of new archetypes, either from the ground up or as specializations of existing ones. Figure 6 shows a possible template for a laboratory report of genomic test results. The top level of the hierarchy is occupied by the composition *Report*, followed by the *Genetic Test Result* observation archetype, developed as a specialization of the existing *Laboratory Test Result*^18^*; to represent actual genetic data we created a new Genetic Findings* cluster, in turn articulated into a number of nested clusters for sequence variation, reference genome and individual variant types (see Table 1).

**Table 1:** Archetypes for genetic data.

| Concept | Type | Status |
| --- | --- | --- |
| Genetic Test Result | OBSERVATION | In progress |
| Genetic Findings | CLUSTER | Created |
| Sequence Variation | CLUSTER | Created |
| Reference Sequence | CLUSTER | Created |
| Substitution Variant | CLUSTER | Created |
| Deletion Variant | CLUSTER | Created |
| Duplication Variant | CLUSTER | Created |
| Insertion Variant | CLUSTER | Created |
| Inversion Variant | CLUSTER | Created |
| Conversion Variant | CLUSTER | Created |
| Indel Variant | CLUSTER | Created |
| Repeated Sequence Variant | CLUSTER | Created |

**Figure 6:**
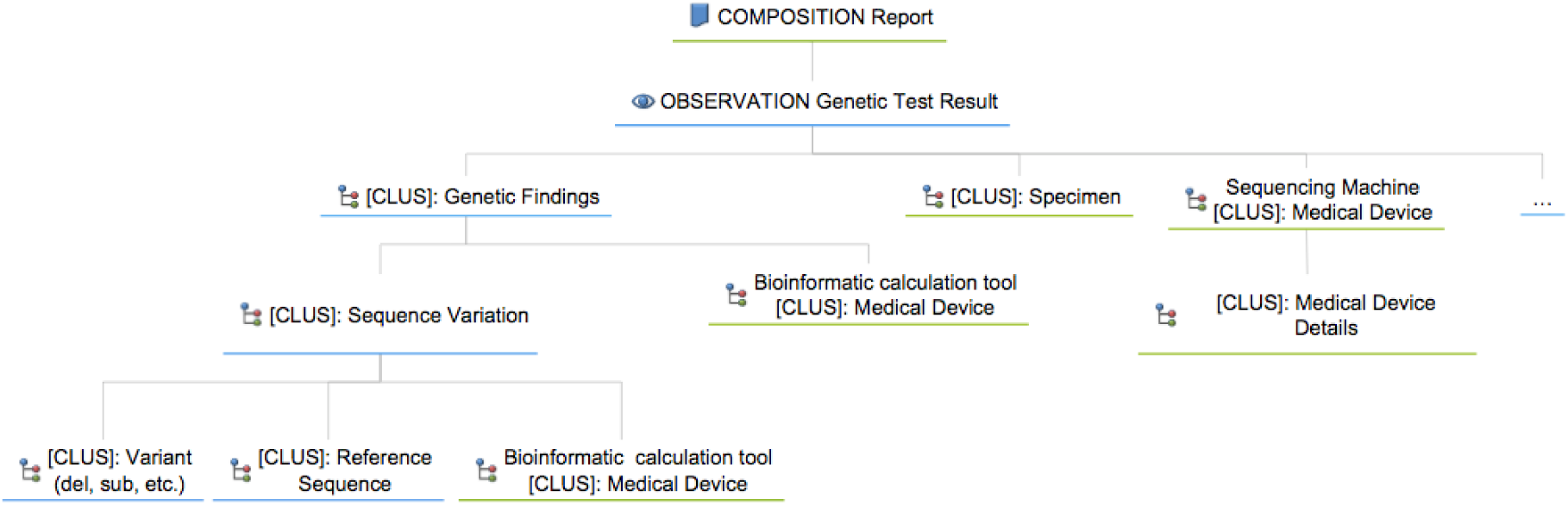
Hierarchical organization of archetypes for genetic content. Existing archetypes are marked with a green border.

### 3.1 OBSERVATION-Genetic Test Result

*Genetic Test Result* models data from a genetic test performed on a patient, along with details of the previously established clinical condition and a description of the protocol. In addition to the attributes carried over from *Laboratory Test Result*, *Genetic Test Result* provides: information on the interpretation and reporting of sequence variations — in accordance with recommendations by the American College of Medical Genetics (ACMG)^19,20^ — in the data section; a representation of the method in the protocol section. The latter, in particular, allows to refer to the bioinformatic workflow as an external resource and to specify the version used to perform the calculation, in order to allow the reconstruction of the data history. With respect to the other archetypes presented here, *Genetic Test Result* is currently at an earlier development stage, and thus more likely to evolve in the near future.

### 3.2 CLUSTER-Genetic Findings

The *Genetic Findings* cluster (see Fig. 8) is meant to be used in the “test findings” slot of *Genetic Test Result*, and can in turn include one or more *Sequence Variation* archetypes to report about variations that are considered relevant for the test. Considering a row in a VCF file, the *Sequence Variation* archetype corresponds to the “standard” parts, while the customizable `INFO` section can be represented by existing fields in *Genetic Finding* or by one or more ad hoc extensions.

**Figure 8:**
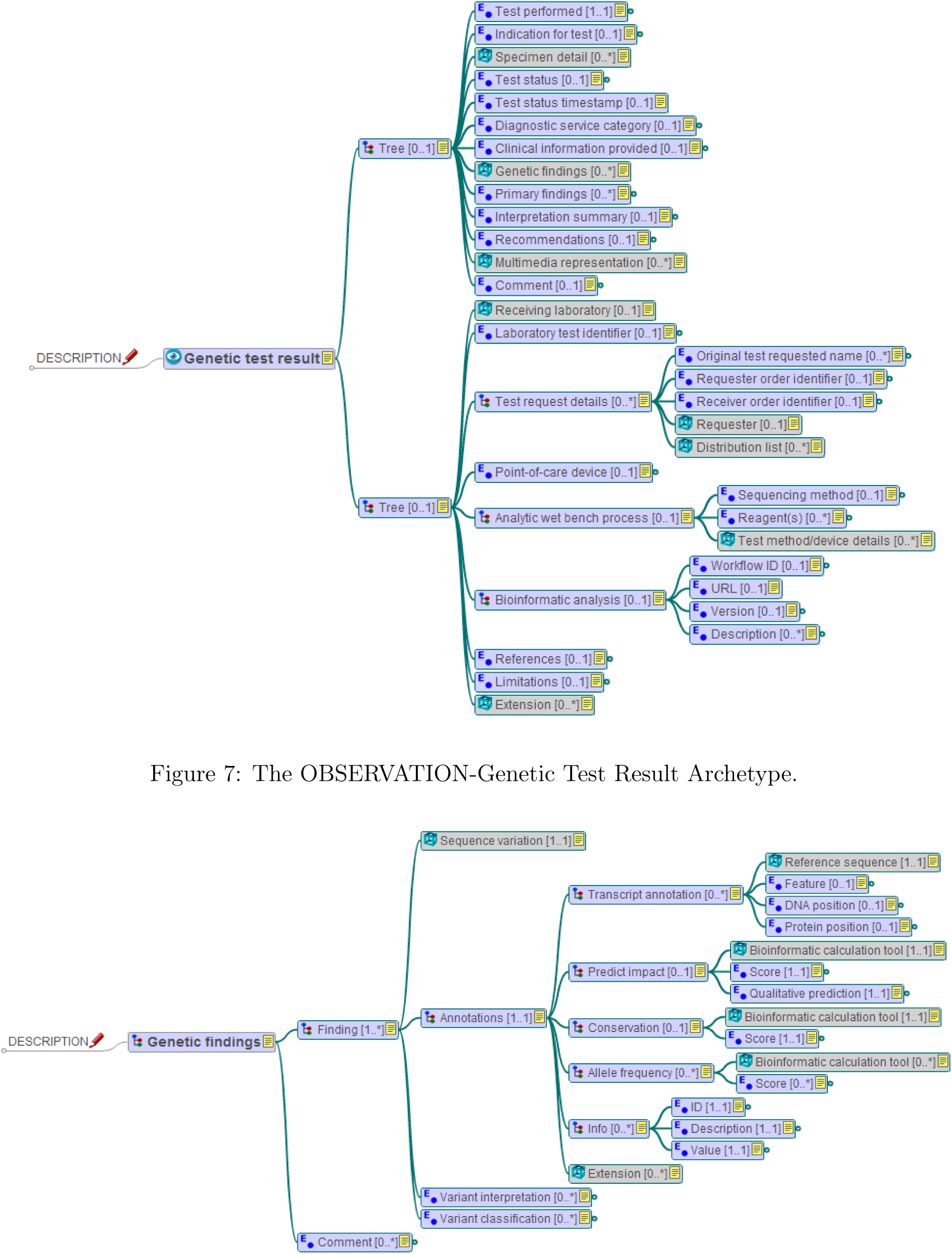
The CLUSTER-Genetic Findings Archetype.

### 3.3 CLUSTER-Sequence Variation and supplementary clusters

The *Sequence Variation* archetype (see Fig. 9) is designed to contain the same data found in a VCF row. The position of the observation in the genome is given with respect to a reference sequence specified in the *Reference Sequence* archetype (Fig. 10). Following the nomenclature proposed by the Human Genome Variation Society (HGVS)^1^, the most common variant types are: substitution, deletion, duplication, insertion, inversion, conversion, insertion-deletion (indel) and repeated sequence^21^. We developed a new cluster archetype for each of the above types (examples are shown in Fig. 11 and 12).

**Figure 9:**
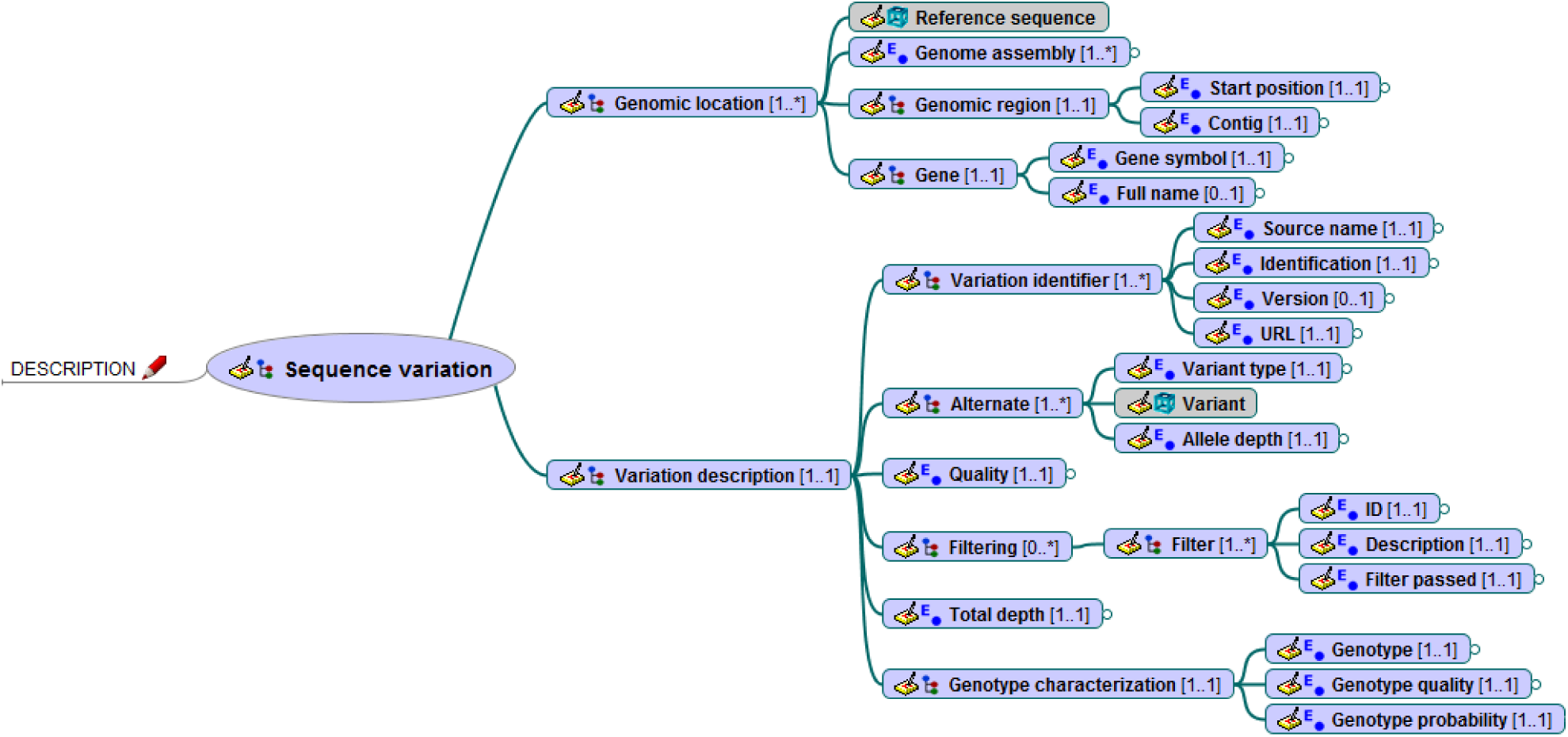
The CLUSTER-Sequence Variation Archetype.

**Figure 10:**
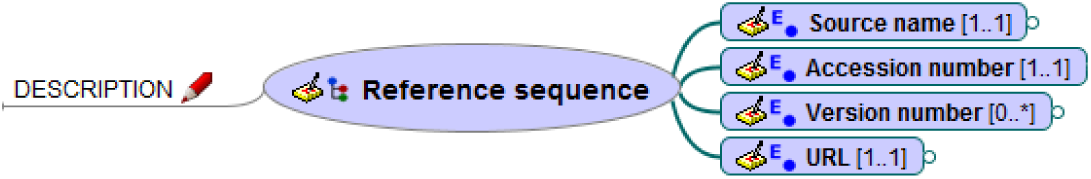
The CLUSTER-Reference Sequence Archetype.

**Figure 11:**
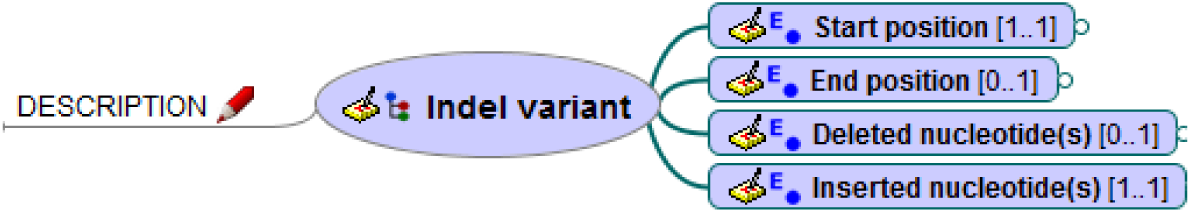
The CLUSTER-Indel Variant Archetype.

**Figure 12:**
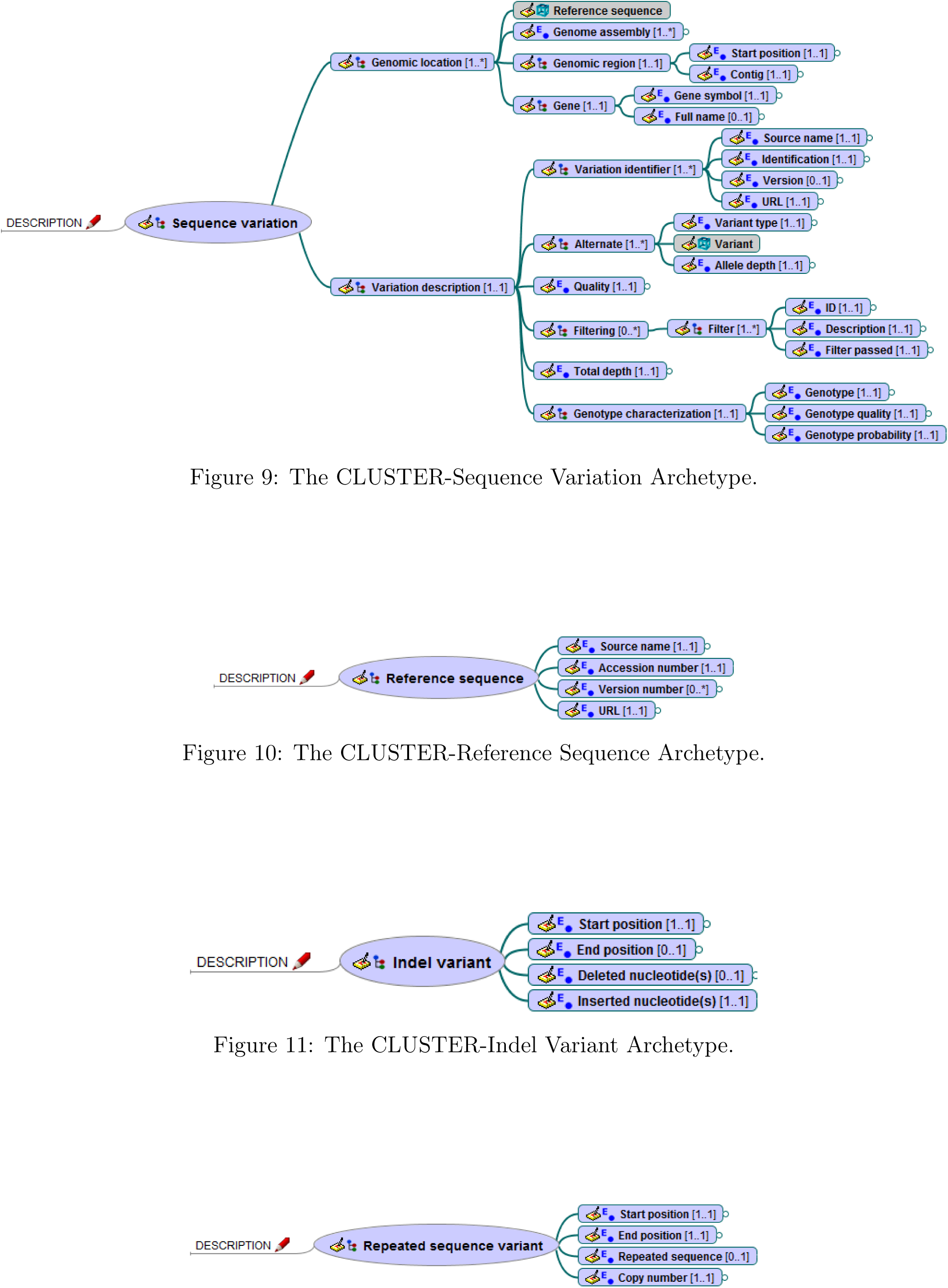
The CLUSTER-Repeated Sequence Variant Archetype.

## 4 Discussion

### 4.1 Real use case application

WES has emerged as a powerful tool for the diagnosis of rare mendelian disorders, especially where standard approaches have failed. This technique focuses on the protein-coding regions of the genome that are more likely to harbour disease-causing variants associated with a particular phenotype. Initially used for research activities, WES is now entering clinics for diagnostic purposes, thanks to the decreasing costs of NGS technologies. A typical study design for the identification of pathogenic variants in rare diseases includes the patient and its parents, and optionally additional family members (affected or not). In this report, we use results related to the WES of one family member to show how openEHR archetypes can successfully describe the genomic information obtained from an NGS-based genetic test (Tables 2 and 5).

**Figure 7:**
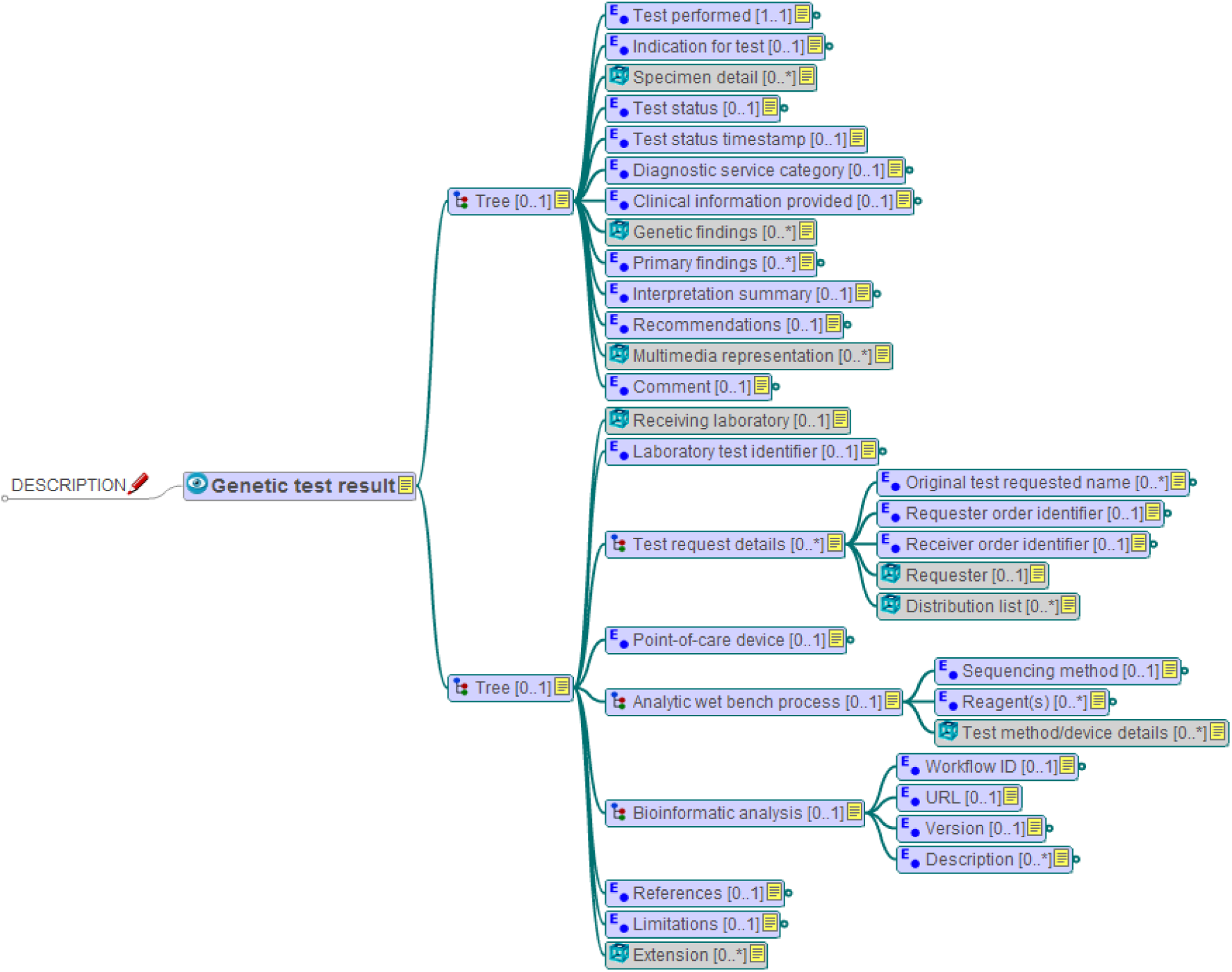
The OBSERVATION-Genetic Test Result Archetype.

**Table 2:**
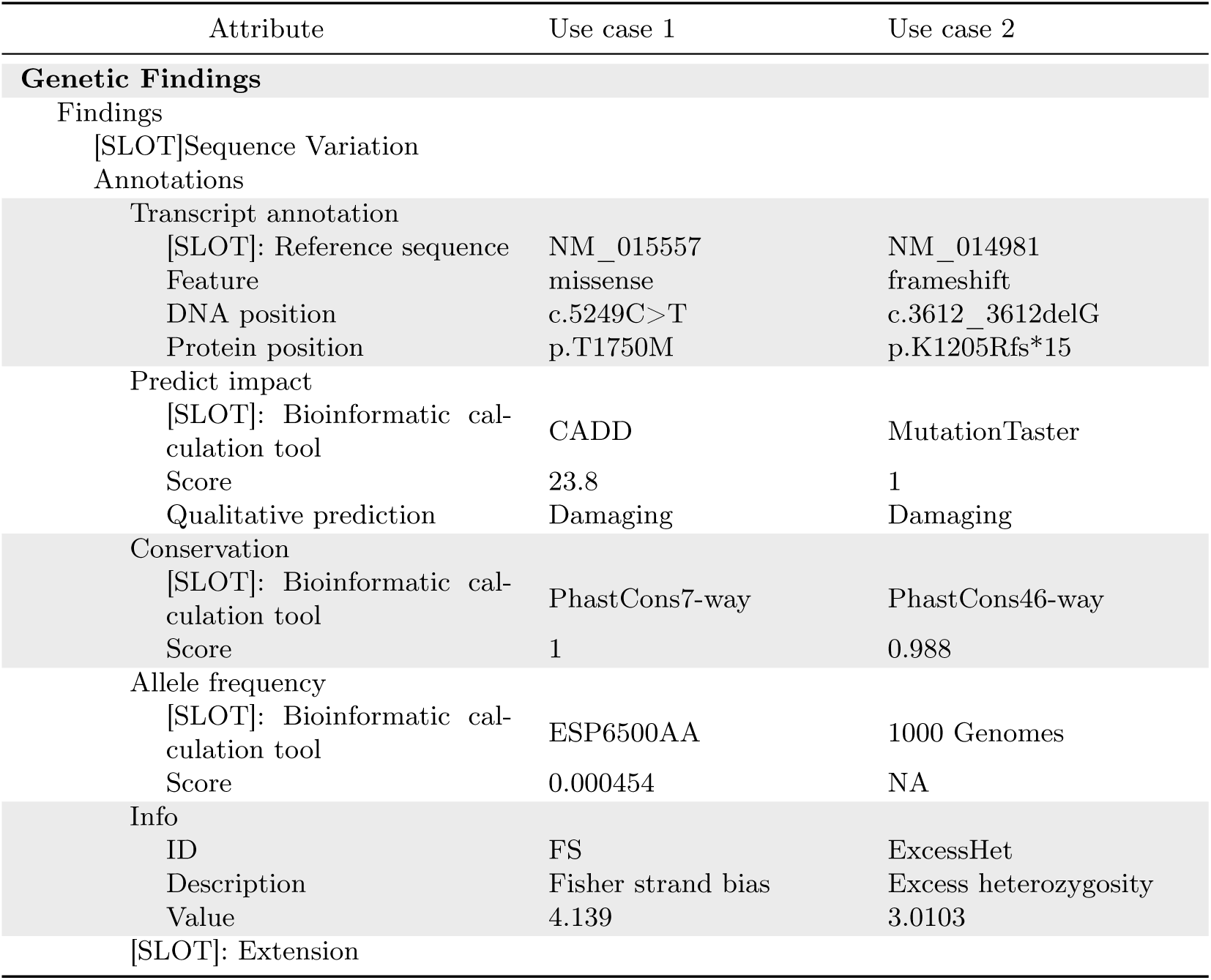
Sample usage of the Genetic Findings archetype

**Table 5:** The comparison was carried out considering three types of data: context, derived and objective.

| Attribute | Use case 1 | Attribute | Use case 2 |
| --- | --- | --- | --- |
| **Substitution Variant** |  | **Deletion Variant** |  |
| Position substituted | 6171835 | Start position | 108147489 |
| Reference nucleotide | G | End position | - |
| New nucleotide | A | Deleted nucleotide(s) | C |

(a) Sample usage of the Substitution Variant archetype
(b) Sample usage of the Deletion Variant archetype

### 4.2 Interoperability: comparison with the HL7^®^ FHIR^®^ standard

This section discusses the interoperability of the proposed model with the one developed by the HL7^®^ Clinical Genomics Work Group as part of the FHIR^®^ standard^22^. HL7^®^ FHIR^®23^ is a standard for exchanging healthcare information that represents granular clinical concepts through basic building blocks called *resources*. Resources are relatively generic and have to be adapted to specific use cases. When a use case is common enough, it may become part of the specification itself as a *profile*. Genomics support in FHIR^®^ consists of four profiles and a resource. The most relevant for this work are the *Observation* profile and the *Sequence* resource. The former, which adapts the *Observation* resource to the genomic context, is used for reporting interpretative genetic information mainly related to variant test results. The latter is instead used to describe an atomic sequence which contains alignment test results and multiple variations. Despite their different reference models, both openEHR and FHIR^®^ can adequately model genetic data, albeit with different degrees of nesting: the openEHR structure is [OBS] Genetic Test Result *→* [CLUS] Genetic Findings *→* [CLUS] Sequence Variation*→* [CLUS] Variant, while the FHIR^®^ one is Observation-Genetics *→* Sequence. The latter, however, can include additional extensions of the *Observation* resource at the same hierarchical level (it grows horizontally, while the openEHR model grows vertically). This has to be taken into account while comparing, together with the different meaning of the “observation” term.

**Table 3:**
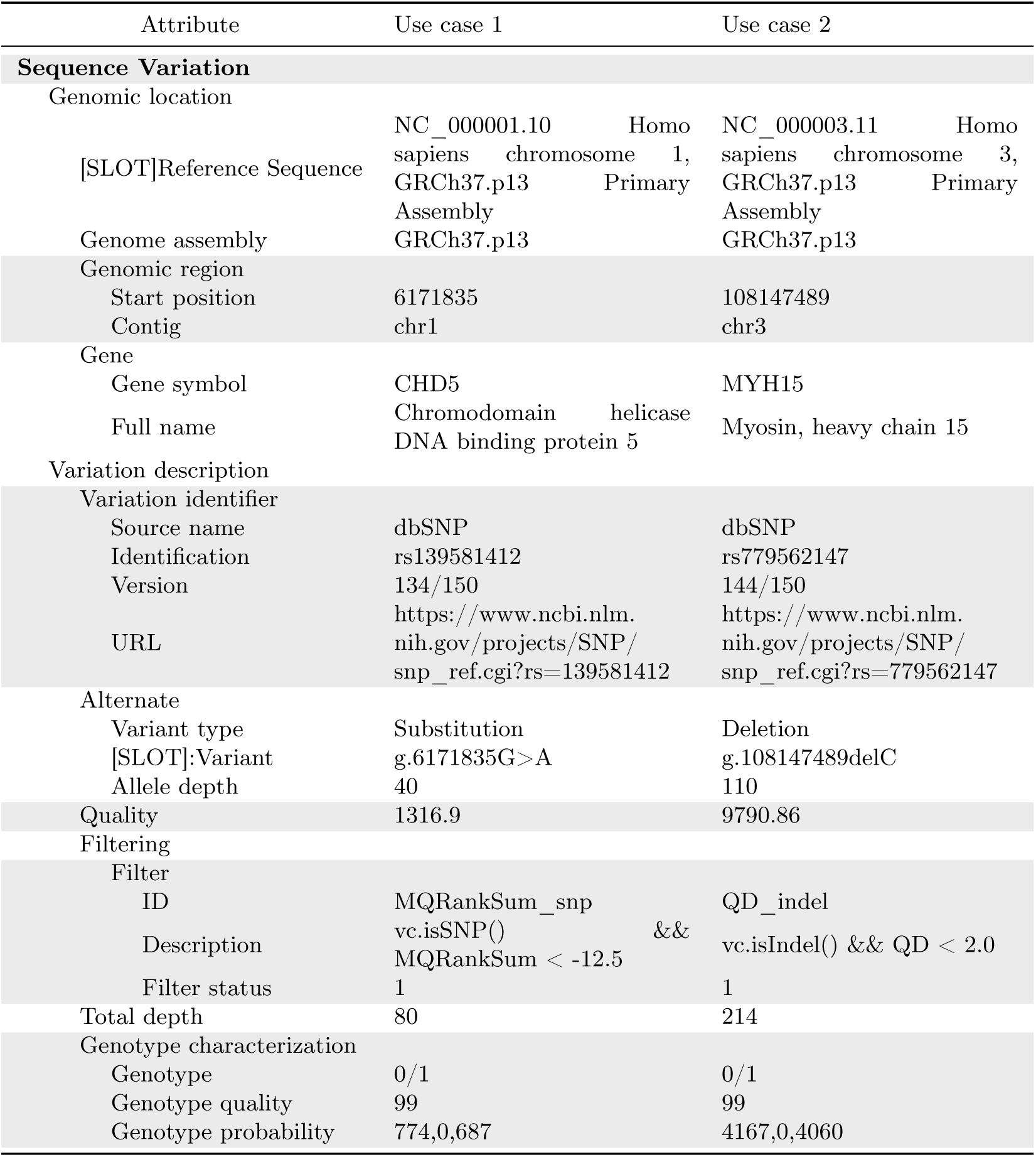
Sample usage of the Sequence Variation archetype

**Table 4:** Sample usage of the Reference Sequence archetype

| Attribute | Use case 1 | Use case 2 |
| --- | --- | --- |
| **Reference Sequence** |  |  |
| Source name | NCBI | NCBI |
| Accession number | NC_000001 | NC_000003 |
| Version number | NC_000001.10 | NC_000003.11 |
| URL | https://www.ncbi.nlm.nih.gov/nuccore/NC_000001.10 | https://www.ncbi.nlm.nih.gov/nuccore/NC_000003.11 |

#### 4.2.1 Context data and interpretations

In the FHIR^®^ model, information about the context of the genetic test performed, the devices used, the specimen, the patient and the performer is given both in the *Observation* resource and in the *Sequence* resource (as a reference to another external resource). Similarly, in the openEHR model the *Genetic Test Result* archetype contains slots to include cluster archetypes (e.g., *Specimen*, *Medical Device*) that provide context information. The difference is that while in FHIR^®^ this information is repeated in multiple places, the recursively nested openEHR approach allows to refer to the same entity multiple times, avoiding redundancy.

Genetic data are typically complemented by subjective information such as diagnosis or generic comments. In the FHIR^®^ model, these are mainly represented within the *Genetics* profile of the *Diagnostic Report* resource. In our model, this type of data could be included in the *Genetic Test Result* archetype.

#### 4.2.2 Derived data

Derived data consist of annotations from the bioinformatic analysis: transcript annotations, impact prediction, sequence conservation, etc. In the openEHR model these items are represented within the *Genetic Findings* cluster archetype, and the range of representable annotations can be broadened via extension slots. At present, FHIR^®^ does not include explicit references to this type of data.

#### 4.2.3 Objective data

Objective data describe the observed sequence variations. In the FHIR^®^ standard, these elements can be found both in the *Sequence* (e.g., reference sequence, variant, quality) and in the *Observation-Genetics* resource (e.g., DNA variant type and ID, gene, allelic state), while in our model they are all included in the *Sequence Variation* archetype, in order to isolate the interpretative aspects from the purely objective variant data.

Our approach differs from the FHIR^®^ one in two main respects: reference sequence and variant description. In FHIR^®^, the reference sequence can be represented by the same *Sequence* resource used to describe the sample sequence, while in our model it is made available via a link to an external entity (repository, accession and version). In FHIR^®^, the observed variant is also represented within the *Sequence* resource through the range of positions affected by the change(s), the reference allele and the observed one and the CIGAR string^15^, while in our model we use a different archetype for each variant type in the HGVS specifications.

### 4.3 Modeling issues

The first challenge encountered during the modeling process was how to represent the results of genetic tests conducted on more than one sample. This happens, for instance, in rare disease studies where patient data is more useful if compared to data from family members, or in oncology when cancer cells are compared to non-cancer cells from the same patient. The archetypes presented here are meant to be used for a single sample, with the assumption that any relationships with other samples are handled at a higher level, i.e., at the composition level.

Another critical aspect concerns the representation of the different types of variant. A simple approach would be to store the reference nucleotide, start position and observed nucleotide(s). This method is very similar to the one adopted in VCF files and works well for single nucleotide substitutions, but can be confusing for more complex variant types. For instance, for simple deletions the VCF `REF` and `ALT` strings must include the base before the event, while this is not required for complex substitutions^16^. To avoid this kind of ambiguity, we used a specific archetype to describe each type of variant.

### 4.4 Related work

Integration of genomic data into EHRs is the subject of several ongoing activities.

The EHR Integration (EHRI) workgroup of the Electronic Medical Records and Genomics (eMERGE) network aims to develop standards and methods for incorporating genomic data into EHRs and optimizing their utilization^24,25^.

The Data Working Group of the Global Alliance for Genomics and Health (GA4GH) concentrates on the representation, storage and analysis of genomic data, with a focus on interoperability^26^. The group developed a web API to allow the exchange of genomic information through a freely available open standard that models entities such as data requests, error messages and actual genomic data fragments^27^.

The Harvard Medical Center for Biomedical Informatics developed the SMART Genomics API^28,29^, an extension of the SMART (Substitutable Medical Applications, Reusable Technologies) platform that uses FHIR^®^ as its base framework to store both Clinical and Genomics data. Following the FHIR^®^ specifications, the team developed new resources and extensions, providing significant input to the HL7^®^ Clinical Genomics Workgroup^30^.

Finally, a first draft of archetypes for genomic data has recently been published in the Norwegian instance of the CKM by the Nasjonal IKT^31^. It consists of five archetypes that describe the patient’s genome, the output of a genetic assessment, the protocol of a genetic laboratory analysis, the description of a variation and of the two alleles. Even though the model is well designed and accurate, it does have a few shortcomings. The *Genetic Lab Analysis* archetype, which describes the protocol used to perform a genetic test, does not support contextual information such as the patient’s clinical conditions. In contrast, we represented genetic test results with a specialization of the existing *Laboratory Test Result* archetype, which contains an accurate description of both context data and the adopted protocol. Moreover, this is in line with openEHR best practice on resources reuse through specialisation^32^. The *Genetic Variant* archetype, together with its *Allele Details* extension, represents information about variations via plain text strings: a characterization (normal or pathogenic), a generic description, one or more reference sequences, etc. In our model, instead, we employ versionable entities (reference sequence, variant ID, etc.) that link to external databases, and we represent each variant type in structured form via a separate cluster archetype. This improves machine readability and makes the archetype more flexible and easily adaptable to the rapid changes in reference sequences and bioinformatic tools. Finally, the IKT model lacks annotations and tools to enable variant filtering and parameter calculation, which we include in the cluster archetype that represents test findings.

### 4.5 Conclusions and future work

We have presented a model for representing genomic content (sequence variation analysis) in a structured form through openEHR archetypes, with the main goals of being machine readable, reusable and shareable. Moreover, each versionable resource or tool involved in the data production process is linked as an external object, allowing to keep track of the particular revision of each instance. We have assessed the feasibility of our approach by applying it to the rare diseases use case. Finally, we have discussed its interoperability with HL7^®^ FHIR^®^.

The archetypes described here are available at https://github.com/crs4/openehr-genomics. In the future, we plan to integrate their use in our production NGS analysis pipelines to formalize the results made available to clinical researchers.

## Footnotes

1 URL: http://varnomen.hgvs.org/

